# Mast Cell Specific *Cyp11a1* Deficiency Promotes T Cell Mediated Immunity and Suppresses Tumour Metastasis in a Mouse Model of Melanoma

**DOI:** 10.64898/2026.03.04.709617

**Authors:** M. Abdul Alim, Katie Butcher, Hosni A.M. Hussein

## Abstract

Mast cells are emerging players in malignant conditions, but the underlying molecular mechanisms remain poorly defined. Based on previous studies showing that steroids can impact on tumour progression in various settings, we here investigated whether mast cell-derived steroid synthesis can have an impact on tumour metastasis in a melanoma model. To this end, we used mice with mast cell-specific ablation of Cyp11a1, a key enzyme in steroid synthesis. We show that lung colonization of melanoma nodules was markedly diminished in mice with mast cell-specific ablation of *Cyp11a1*, accompanied by reduced infiltration of mast cells into the lungs. *Cyp11a1* gene expression was significantly decreased in lungs of mice with mast cell-specific ablation of Cyp11a1, indicating that mast cells account for a substantial fraction of the total Cyp11a1 expression. Our results also revealed that the mast cell-specific deletion of *Cyp11a1* led to an overall increase in CD107a/LAMP1 staining intensity of the lung tissue, suggesting that mast cell-derived steroids can suppress immune cell activation and degranulation. A further dissection of this finding by flow cytometry analysis of individual immune cell populations revealed that CD8^+^ T cells, NK cells and basophils were activated to a higher extent in lungs from mice with mast cell-specific *Cyp11a1* ablation. We also demonstrate that both CD8^+^ and CD4^+^ T cells in lungs of mice with mast cell-specific deletion of *Cyp11a1* expressed elevated levels of IFN-γ in comparison with controls. Altogether, these findings introduce a hitherto unrecognized role of a mast cell-derived steroid axis in regulating tumour metastasis.

## INTRODUCTION

Although being traditionally recognized for their role in allergic responses and pathogen defence, mast cells are increasingly recognized as active contributors in the tumour microenvironment, where they impact disease progression through a range of immunological and structural mechanisms [1-3]. Mast cells are key players in neuroimmune communication, mediating interactions with the peripheral nervous system and the immune system. A role for mast cells has also been suggested in range of additional settings, including tissue healing responses and inflammation, and metabolic disorders such as diabetes, hyperlipidemia and thyroiditis [4-8]. Given their known ability to interact with other immune cells and their capacity to influence extracellular matrix parameters, it is conceivable that mast cells can have an impact on tumour development and metastasis. However, this issue has not been studied to a large extent, and there is thus a need for a deeper understanding of mast cell functions in cancer immunology. A number of studies have approached these issues, and have revealed that mast cells can both suppress and promote tumourigenesis depending on the context [9, 10]. By releasing a wide range of preformed and newly synthesized mediators including cytokines (e.g., TNF□α, IL□6, IL□10, IL-33), proteases (e.g., tryptase, chymase), lipid mediators (e.g., prostaglandins, leukotrienes), and growth factors such as VEGF, mast cells may have the capacity to shape angiogenesis, immune recruitment, and extracellular matrix dynamics [11-13]. However, the precise molecular pathways through which mast cells influence tumour progression and metastatic niche formation remain poorly understood [10, 14, 15].

De novo steroidogenesis has emerged as a non-classical immunomodulatory mechanism. In this process, the mitochondrial rate limiting enzyme CYP11A1 catalyses the conversion of cholesterol into pregnenolone - a reaction that initiates steroid hormone biosynthesis [16, 17]. Steroid biosynthesis, though classically attributed to endocrine tissues, has increasingly been documented in immune populations, where locally synthesized steroids can act as immunoregulatory signals [18-21]. Further, it has recently been shown that tumour-derived pregnenolone promotes bone metastasis by enhancing osteoclastogenesis through prolyl 4□hydroxylase subunit beta (P4HB), in breast cancer and melanoma models [22]. It has also been demonstrated that tumour-associated immune cells can utilize the steroidogenesis pathway to promote immune suppression [23, 24]. However, the role of mast cell steroidogenesis and its relevance in shaping the immune landscape within tumour contexts has not been explored.

In this study, we investigated the role of mast cell-derived steroidogenesis in shaping the immune microenvironment during pulmonary metastasis of melanoma. Using a genetically engineered mouse model with mast cell-specific ablation of Cyp11a1, we demonstrate that mast cell–derived steroidogenesis mediates immune suppression within lung tumours, which is associated with enhanced lung tumour colonization. Hence, this study introduces a novel mast cell-steroid axis in regulating tumour progression.

## MATERIALS AND METHODS

### Mice and Ethical Approval

In this study, we used C57BL/6J mice, originally sourced from The Jackson Laboratory. Mast cell-specific Cyp11a1 knockout mice (cKO) were generated by crossing Cpa3-Cre transgenic mice (JAX stock #026828, Jackson laboratory) with Cyp11a1^fl/fl^ animals (generated by Sanger Institute) [25, 26]. Littermates possessing the Cyp11a1^fl/fl^ genotype but lacking the Cpa3-Cre transgene served as wild-type (WT) controls. All mice were housed in a specific pathogen-free (SPF) facility under controlled conditions, including a 12-hour light/12-hour dark cycle. Both male and female mice were included, with random allocation to experimental groups to minimize bias. All procedures involving animals adhered to the UK Animals (Scientific Procedures) Act 1986 and were conducted under Home Office Project License (PPL PP2448972). Ethical approval was obtained from the institutional Animal Welfare and Ethical Review Body, described elsewhere [26].

### Melanoma Lung Metastasis Model

For metastatic challenge, 0.5 × 10^6^ B16-F10 melanoma cells were injected intravenously via the tail vein into 12-18-weeks-old control (n=8) and cKO (n=8) mice. Animals were euthanized at day 13 post-injection for endpoint analyses unless otherwise indicated (**Fig. 1**). To assess general health and systemic effects of tumour burden, body weight was recorded at baseline (day 0), day 7, and day 13 following tumour cell injection. Weight measurements were performed at the same time each day using a digital scale. The mice weight was 19.9-27.7 g for WT controls, and 23.8-34.6 g for cKO mice, and were aged between 10-18 weeks at the start of the experiment. At the conclusion of the study, all mice were humanely euthanized by cervical dislocation in accordance with institutional guidelines, ensuring a rapid and ethical procedure.

**Figure 1.**
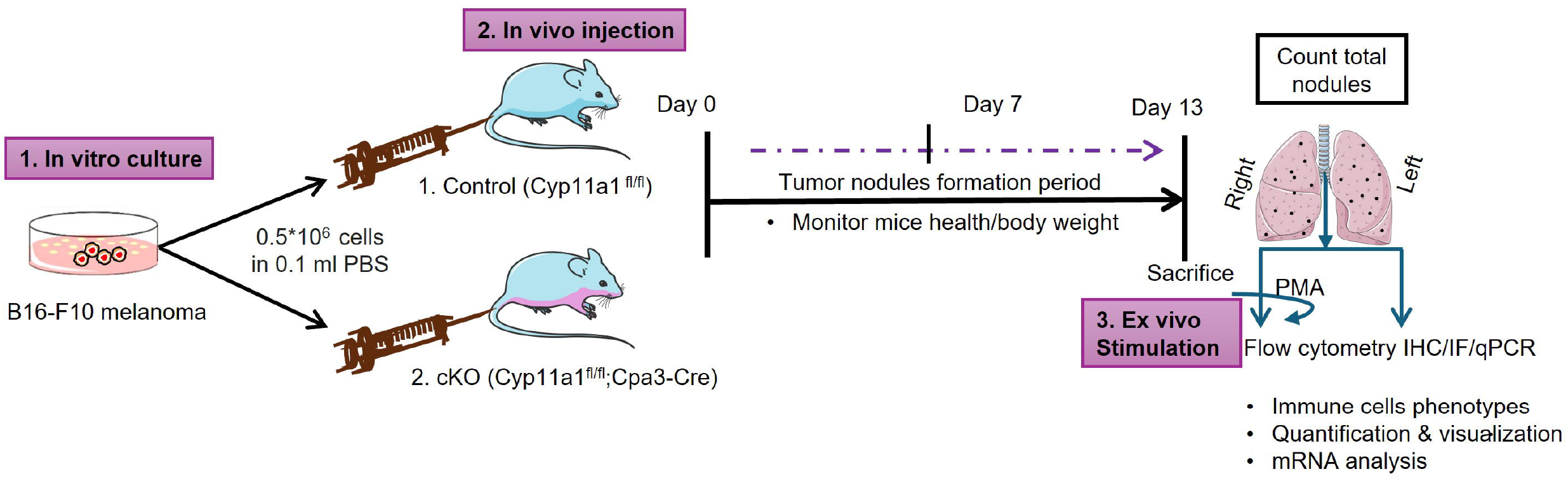
Schematic illustration of the experimental design for the melanoma lung metastasis model. Control and conditional knockout (cKO) mice were intravenously injected with 0.5 × 10□ B16-F10 melanoma cells, and assessed for pulmonary tumour nodule formation at day 13. Body weight was monitored on days 0, 7, and 13. Mice were euthanized on day 13, and total tumour nodules were counted. Next, lung tissue samples were processed for characterization of immune cells, following ex vivo stimulation with PMA.

### Tumour Burden Quantification and Tissue Processing

At endpoint, lungs were perfused by ice-cold PBS and harvested in PBS supplemented with 2.5% FBS and 2 mM EDTA and stored on ice until further processing. Lungs were rinsed in ice-cold PBS to enhance nodule visibility. Visible surface tumour nodules were counted manually with two independent observers (blinded) as well as under a microscope after basic histological staining, and representative images were captured and quantified in a semi-quantitative fashion.

For histological analysis, a small portion of lung tissue was immediately frozen (-80°C). The remaining tissue was placed in PBS within a 2□ml microcentrifuge tube and manually chopped into small pieces using sterile scissors. A digestion cocktail was freshly prepared with DMEM/F12 supplemented with125□μl (1□mg/ml) of collagenase A (Roche), 125□μl (1□mg/ml) of collagenase D (Roche), and 35 μl of DNase I (0.2–0.4□mg/ml). Samples were incubated at 37°C with gentle agitation (Eppendorf ThermoMixer C) for 30 minutes. To terminate enzymatic activity, cells were washed in PBS containing 2% FBS and 2 mM EDTA (FACS buffer). Digested tissue was filtered through 70□μm nylon mesh strainers (Falcon) into 50□ml conical tubes and washed with 30□ml cold PBS. Cell suspensions were centrifuged at 400× g for 5 min at 4°C. Pellets were briefly vortexed and incubated with 3□ml of 1× red blood cell lysis buffer at room temperature for 3 min. The lysis reaction was halted by dilution with PBS, followed by centrifugation under the same conditions. Cells were resuspended in PBS, filtered through 30□μm strainers, and kept on ice for immediate downstream processing.

### Histology and Toluidine Blue Staining

Frozen lung tissues were sectioned at 7 μm thickness under a cryostat (-20°C) and the slices were fixed with ice-cold acetone (-20°C) for 20□min. Sections were stained with hematoxylin and eosin (H&E) to evaluate tumour architecture. For mast cell visualization, adjacent sections were stained with Toluidine blue (0.1% in 0.5N HCl) for 3 min at room temperature. Mast cells were quantified in peritumoural regions from at least three-five high-power fields (HPF) per section. Staining was imaged using a Zeiss AxioObserver.Z1 Widefield microscope.

### Immunofluorescence and Confocal Microscopy

Lung tissue slices from control (Cyp11a1-sufficient) and mast cell–specific Cyp11a1 knockout (cKO i.e., Cyp11a1-deficient) mice were fixed in cold acetone (-20□°C) for 20 min. Staining with antibodies was performed following protocols described previously [4, 27] Briefly, tissue sections were blocked with normal horse and/or goat serum, followed by three washes in PBS. Slides were then incubated overnight at 4°C in the dark with the following primary antibodies: Monoclonal Mouse Mast Cell Tryptase (ab2378, Abcam) diluted 1:100, CYP11A1 (D8F4F) Rabbit mAb (Cell Signaling Technology, #14217) diluted 1:200, Mouse CD45 (FITC) diluted 1:200, and CD107a PE diluted 1:200.

After incubation with primary antibodies, tissue sections were washed three times with PBS and subsequently incubated for 60 min at room temperature with secondary antibodies. These included anti-rabbit IgG F(ab’)□ Alexa Fluor 488 (Cell Signaling Technology, #4412S) or goat anti-rabbit multi-rAb CoraLite Plus 555 (Proteintech, #RGAR003), both diluted 1:200, and F(ab’)□ goat anti-mouse IgG Alexa Fluor 647 (Proteintech, #SA00014-10), also diluted 1:200. Nuclei were counterstained with DAPI (4′,6-diamidino-2-phenylindole; Invitrogen).

Confocal microscopy was performed using a Zeiss LSM700 confocal microscope. Images were captured at original magnifications of ×200 or ×400. Image processing and analysis were conducted using ImageJ software (Fiji, ImageJ2).

### Flow Cytometry

The resulting single-cell suspensions were stained for flowcytometry using FACS buffer (PBS containing 2% FBS and 2 mM EDTA). Cells were incubated with anti-CD16/32 (Fc block; eBioscience or BioLegend) for 10□min at 4°C, followed by surface staining in FACS buffer with fluorochrome-conjugated antibodies against the following markers: CD45 (1:1,000), CD11c or I-A/I-E (1:1,000), CD4 (1:1,000), CD8 (1:400), CD11b, FcεRI (1:400), NK1.1 (CD161) (1:500), CD117 (c-Kit), CD170 (Siglec-F), CD19 (1:800), and CD107a (1:200). Dead cells were excluded using Ghost Dye UV 450 (CYTEK, 1:1,000).

For intracellular cytokine staining, cells were stimulated overnight at 37°C with PMA (50□ng/mL) and ionomycin (500□ng/mL) in the presence of brefeldin A (5□μg/mL; BD Biosciences). After surface staining and fixation/permeabilization (BD Cytofix/Cytoperm kit), cells were stained with antibodies against CD3 (1:800), IFN-γ (1:400), and TNF-α (1:400). All antibodies were titrated for optimal performance. Data acquisition was performed on a Cytek Aurora spectral flow cytometer (Cytek Biosciences), and spectral unmixing and data analysis were conducted using FlowJo v10. Immune cell subsets were quantified as a percentage of live CD45□ cells. Mast cells were identified as FcεRI□ CD117□.

### RNA Extraction, cDNA Synthesis and Quantitative PCR

Frozen lung tissue samples were homogenized, and total RNA was extracted using the RNeasy Mini Kit (Qiagen) following the manufacturer’s guidelines. RNA quantity was determined using a NanoDrop spectrophotometer (Thermo Fisher Scientific), and RNA integrity was assessed with the Agilent 2100 Bioanalyzer (Agilent Technologies).

For cDNA synthesis, 250 ng of total RNA was reverse transcribed using the iScript cDNA Synthesis Kit (Bio-Rad), according to the provided protocol. Quantitative PCR was carried out using SYBR Green Supermix (Bio-Rad) on a CFX384 Real-Time PCR System (Bio-Rad). Primer sequences were either designed using PrimerBank or obtained as pre-validated primer assays from Bio-Rad (Supplementary Table 1). All primer pairs were tested for amplification efficiency prior to use to ensure accurate quantification.

Relative gene expression levels were calculated using the comparative Ct (ΔΔCt) method. Actb and Gapdh were used as endogenous reference genes for normalization. All reactions were run in technical duplicates. Statistical analysis was performed using unpaired two-tailed t-tests to assess differences in gene expression between groups.

### Statistical analyses

Data are presented as mean□±□standard error of the mean (SEM). Comparisons between two groups were made using unpaired, two-tailed Student’s t-test. P values < 0.05 were considered statistically significant. All analyses were performed using GraphPad Prism (v10.0). However, due to unequal group sizes of mice (M/F) and the study was not being powered to detect sex-specific differences, data from male and female mice were pooled for all analyses. Sex was therefore not included as an independent variable in statistical comparisons.

## RESULTS

### Mast cell-derived Cyp11a1 promotes lung metastatic colonization of melanoma cells and mast cell infiltration

To investigate the role of mast cell-expressed Cyp11a1 in metastatic tumour growth, we compared lung tumour burden in control (Cyp11a1-sufficient) and mast cell-specific *Cyp11a1* knockout (cKO) mice following intravenous injection of melanoma cells (**Fig. 1**). Control mice exhibited a significantly greater number of lung tumour nodules relative to cKO counterparts, indicating that mast cell-derived Cyp11a1 facilitates metastatic colonization and tumour outgrowth in the lung microenvironment (**Fig. 2a-b**). Importantly, body weight monitoring over the course of tumour progression revealed no significant differences between groups, excluding systemic illness or cachexia as confounding factors for tumour burden disparities (**Fig. 2c**). Histological analysis of lung sections using hematoxylin/eosin and toluidine blue staining demonstrated enhanced tumour nodule formation and increased mast cell infiltration in control lungs compared to those from cKO mice (**Fig. 2d-e**). In agreement with this, quantitative real time PCR (qPCR) measurements demonstrated that the expression of mast cell-specific genes (Mcpt1, Mcpt6) was profoundly diminished in lungs of cKO vs control mice (**Fig. 2f-g**). These findings indicate that mast cell-expressed Cyp11a1 promotes melanoma colonization of the lung, and that mast cell-expressed Cyp11a1 has an autocrine function to support mast cell accumulation in the context of melanoma metastasis to the lung.

**Figure 2.**
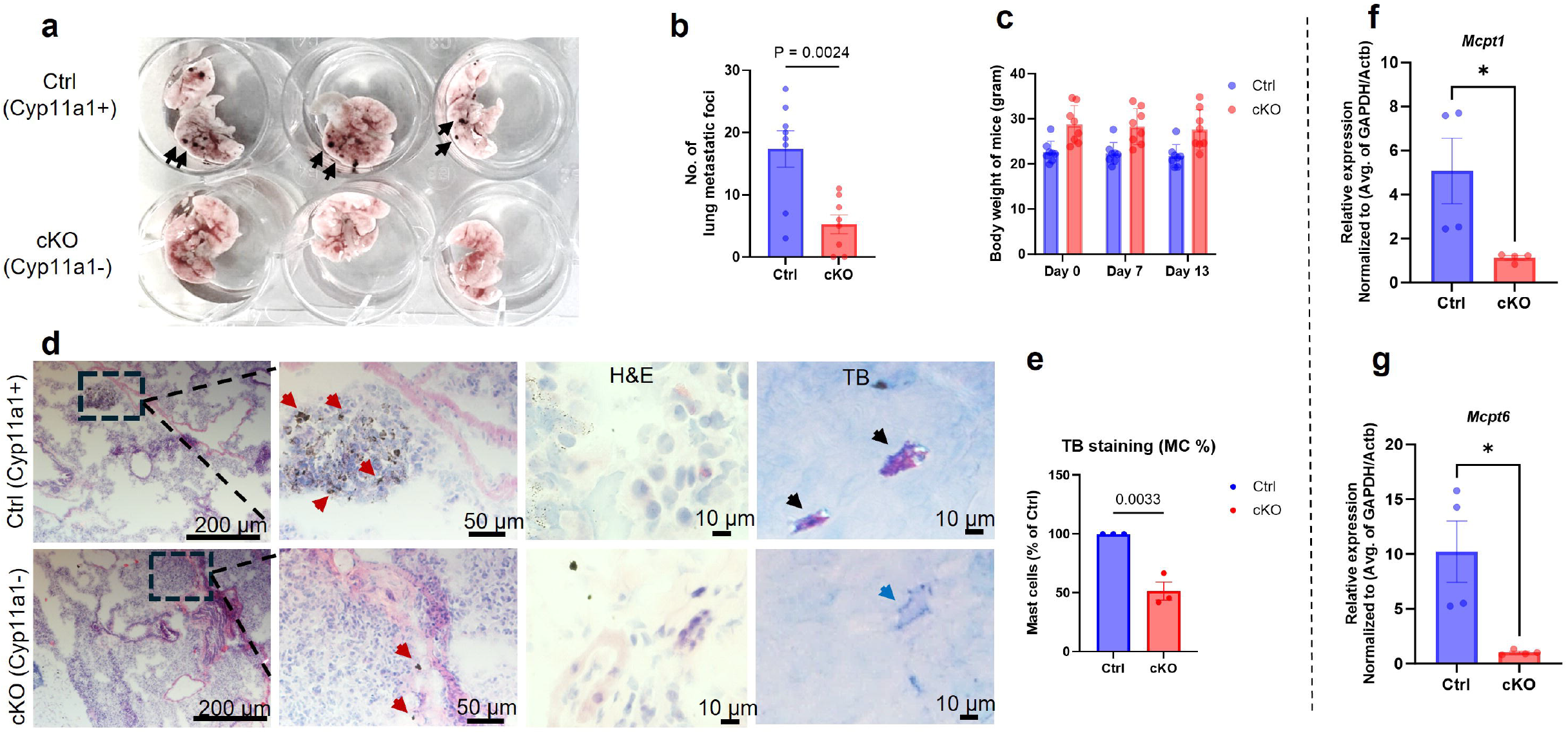
Mast cell–derived Cyp11a1 promotes lung tumour colonization and is associated with increased mast cell infiltration. (**a**) Representative images of lung tumour nodules from control (Cyp11a1-sufficient) and mast cell–specific *Cyp11a1* knockout (cKO) mice following metastatic challenge. Note the greater tumour burden (black arrows) in control vs. cKO animals. (**b**) Quantification of lung metastatic nodules. (**c**) Body weight monitoring of mice during metastatic progression. (**d**) Histological (H&E) and toluidine blue staining showing tumour tissue morphology with enhanced nodule formation (red arrows) and greater mast cell infiltration (black arrows) in lung tissues from control (Cyp11a1-sufficient) mice compared to mast cell–specific *Cyp11a1* knockout mice. (**e**) Quantitative assessment of mast cells in lungs of control- and cKO mice (at least three-five high-power fields per section were counted). **(f-g)** Quantitative RT-PCR (qPCR) analysis of the expression of mast cell marker genes (*Mcpt1* and *Mcpt6*) in lungs of tumour-bearing control (Ctrl) and cKO mice. Gene expression was normalized to *GAPDH* and *Actb* using the ΔΔCt method and presented as fold change. Statistical analysis was performed using unpaired *t*-tests (For tumour nodule enumeration, n = 8; histology, n = 3; qPCR, n = 4). *P* < 0.05 was considered significant.

### Cyp11a1 is expressed in the mast cell niche in lung tumours

Confocal microscopy analysis of lung tumour sections revealed a large extent of colocalization of tryptase (a mast cell marker) and the steroidogenic enzyme Cyp11a1 in control (Cyp11a1-sufficient) mice, indicating a high abundance of steroidogenic mast cells within the metastatic lung microenvironment (**Fig. 3a-b**). In contrast, tumours from mast cell-specific Cyp11a1 knockout (cKO) mice exhibited markedly reduced Cyp11a1: tryptase colocalization, consistent with diminished Cyp11a1 expression in mast cells (**Fig. 3a-b**).

**Figure 3.**
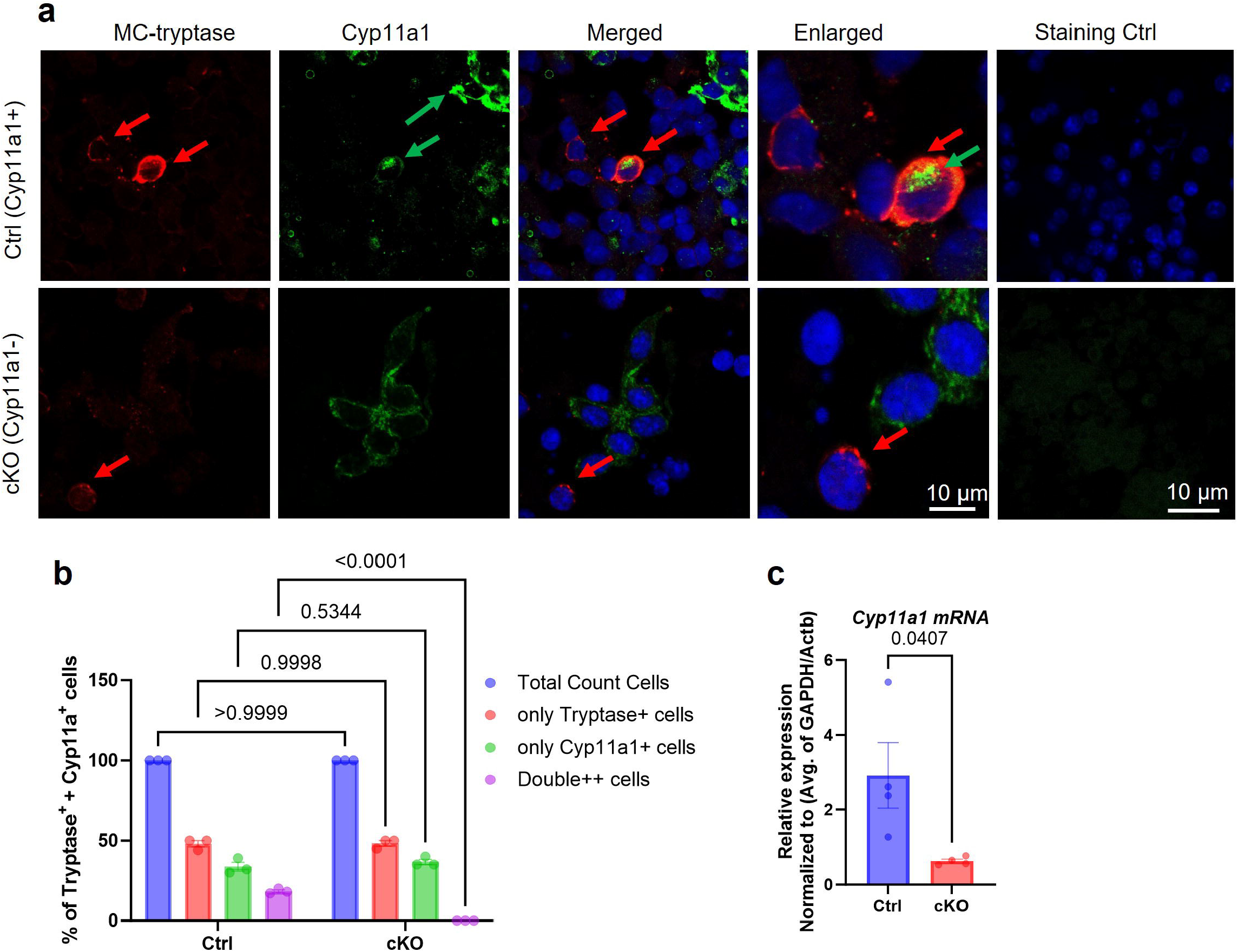
Expression of Cyp11a1 in mast cells of tumours from control mice but not in mice with mast cell-specific depletion of *Cyp11a1*. (**a**) Confocal microscopy images showing localization of the mast cell marker tryptase (red) and Cyp11a1 (green) in lung tumours from control (Ctrl; Cyp11a1-sufficient) and mast cell–specific Cyp11a1 knockout (cKO) mice. (**b**) Quantitative analysis of Cyp11a1/tryptase co-localization using ImageJ software confirming a significantly higher proportion of Cyp11a1□ mast cells in control tumours compared to cKO tumours (**c**) Quantitative RT-PCR (qPCR) analysis of *Cyp11a1* gene expression in lungs of tumour-bearing control (Ctrl) and cKO mice. *Cyp11a1* gene expression was normalized to *Gapdh* and *Actb* expression using the ΔΔCt method and presented as fold change. Statistical analysis was performed using unpaired t-tests (For IF staining, n = 3; for qPCR, n= 4). P < 0.05 was considered significant.

### Targeted deletion of *Cyp11a1* in Mast cells is associated with reduced CD45□ immune cell activation in lung tumours

Next, we asked whether mast cell-specific deletion of Cyp11a1 has an effect on the overall population of Cyp11a1-positive cells, and on the overall accumulation of leukocytes in the lung tumours. To this end, tumour sections were stained for the pan-leukocyte marker CD45 (red) and Cyp11a1(green) (**Fig. 4a**). These analyses revealed that the overall intensity of Cyp11a1 staining was markedly diminished in tumours from cKO vs control mice, indicating that mast cells account for a large fraction of the total population of Cyp11a1-positive cells. The reduction of overall Cyp11a1 expression in lungs of cKO mice was also supported by qPCR analysis, which revealed a profound reduction in *Cyp11a1* mRNA levels in lungs of cKO vs control mice (**Fig. 3c**).

**Figure 4.**
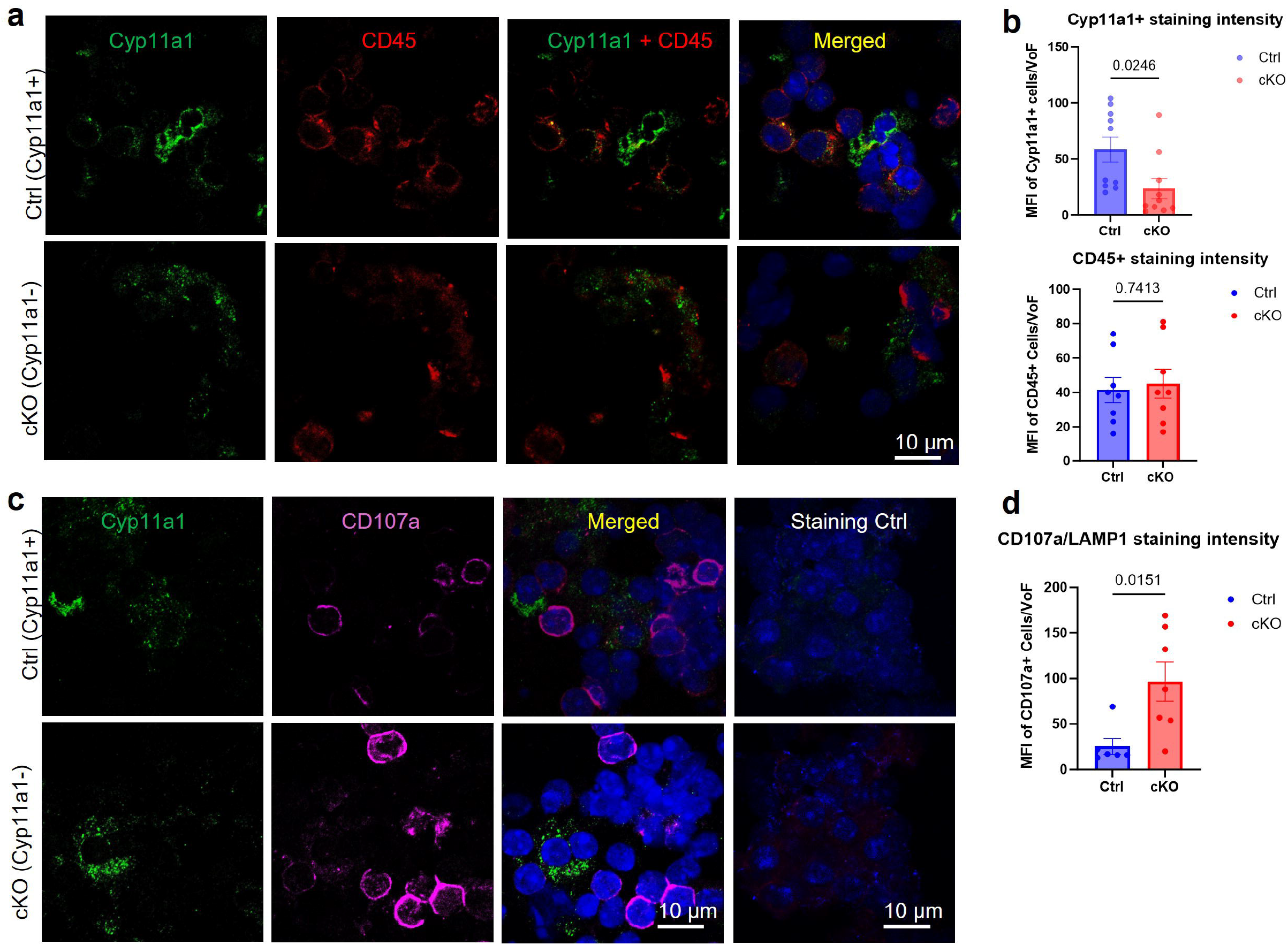
Expression of Cyp11a1, CD45 and CD107a in lung cells from tumour-bearing control mice and mice with Cyp11a1-deficient mast cells. (**a, c**) Confocal microscopy images showing expression of the general leukocyte marker CD45 (red), the activation marker CD107a (magenta) and Cyp11a1 in lung cells isolated from control (Cyp11a1-sufficient) and mast cell–specific *Cyp11a1* knockout (cKO) tumour-bearing mice. (**b, d**) Quantitative analysis of the intensity of Cyp11a1, CD45 and CD107a staining intensity in lung tissue cells from control mice (Ctrl) and mice with mast cell-specific depletion of *Cyp11a1* (cKO), using the ImageJ software. VoF, view of field. Data represent mean values ±□SEM; *P*□< □0.05 by unpaired two-tailed *t*-test (n = 3).

In contrast, similar intensities of CD45 staining were seen in tumours of both control- and cKO mice (**Fig. 4b**; lower panel), suggesting that the absence of Cyp11a1 expression in mast cells does not affect the overall immune cell infiltration to the lung. To assess for the extent of overall immune cell degranulation, we stained for CD107a (also known as LAMP1), a protein that is transported to the cell surface upon degranulation of immune cells (including mast cells, cytotoxic T lymphocytes, basophils, eosinophils). Interestingly, the overall positivity for CD107a was markedly higher in cKO vs control mice (**Fig. 4c-d**). indicating that mast cell-dependent steroidogenesis regulates immune cell activation within the lung tumours.

### Effect of mast cell–derived Cyp11a1 on lung immune cell populations

To provide further insight into how mast cell-dependent steroidogenesis regulates the immune landscape of the melanoma tumours of the lung, we performed flow cytometry analysis of various immune cells subsets. These analyses demonstrated comparable frequencies of T cells (CD4^+^ and CD8^+^), B cells, NK cells, eosinophils and basophils between control- and mast cell-specific *Cyp11a1* knockout (cKO) mice during baseline conditions (**Fig. 5**).

**Figure 5.**
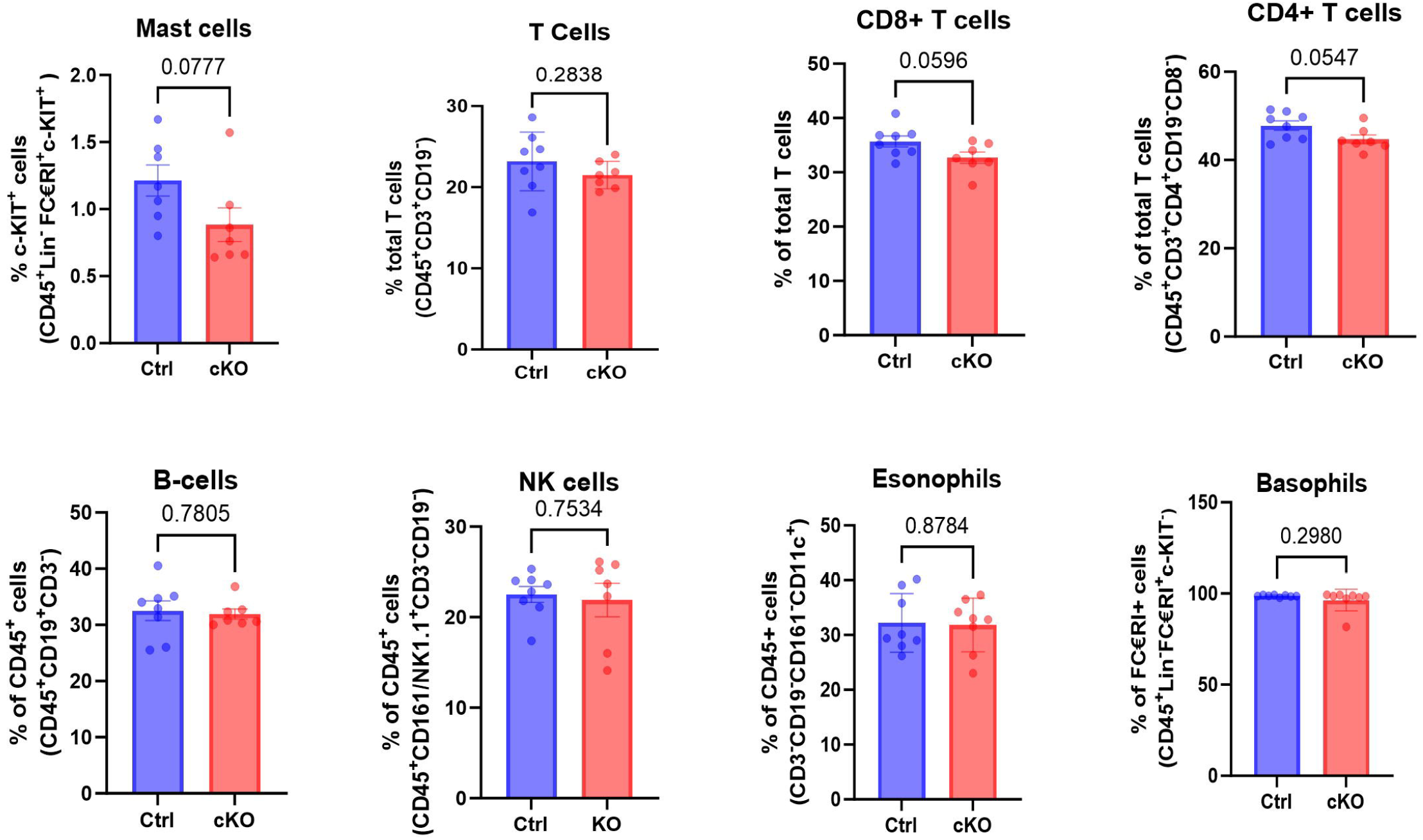
Immune profiling of lungs from tumour-bearing control mice and mice with Cyp11a1-deficient mast cells. Immunophenotyping of lung tumours by flow cytometry. Note that no significant effect of mast cell-specific depletion of *Cyp11a1* on the proportion of T cells, B cells, NK cells, eosinophils or basophils among the CD45^+^ population were seen (gating strategy shown in **Suppl. Fig. 1**). Data represent meanLvalues ±□SEM; *P*□<□0.05 by unpaired two-tailed *t*-test (Ctrl, n = 8; cKO, n = 7 - 8).

### Loss of mast cell-derived Cyp11a1 enhances immune effector cell activation

Based on the demonstration that mast cell-specific expression of Cyp11a1 suppresses the overall population of CD107a-positive (degranulated) immune cells in the lung melanoma tumours, we next assessed corresponding effects on individual immune cell types. As seen in **Fig. 6**, the proportions of activated (degranulated) CD8^+^ T-cells, NK cells and basophils were significantly higher in the tumours from cKO vs control mice at baseline (unstimulated; US), hence indicating that mast cells, via Cyp11a1, suppressed the activation of these immune cell populations. After PMA stimulation (S), the expression of CD107a remained significantly higher in CD8^+^ T cells and basophils in the lungs of cKO vs control mice, whereas CD107a expression in NK cells was not significantly different between the genotypes after PMA stimulation (**Fig. 6**). Notably, although the depletion of mast cell-expressed *Cyp11a1* did not affect the cell surface expression of CD107a in mast cells at baseline conditions, we noted that PMA-activated cKO mast cells exposed lower levels of cell surface CD107a than control cells, possibly reflecting an effect of intrinsic steroidogenesis on their capacity to undergo activation.

**Figure 6.**
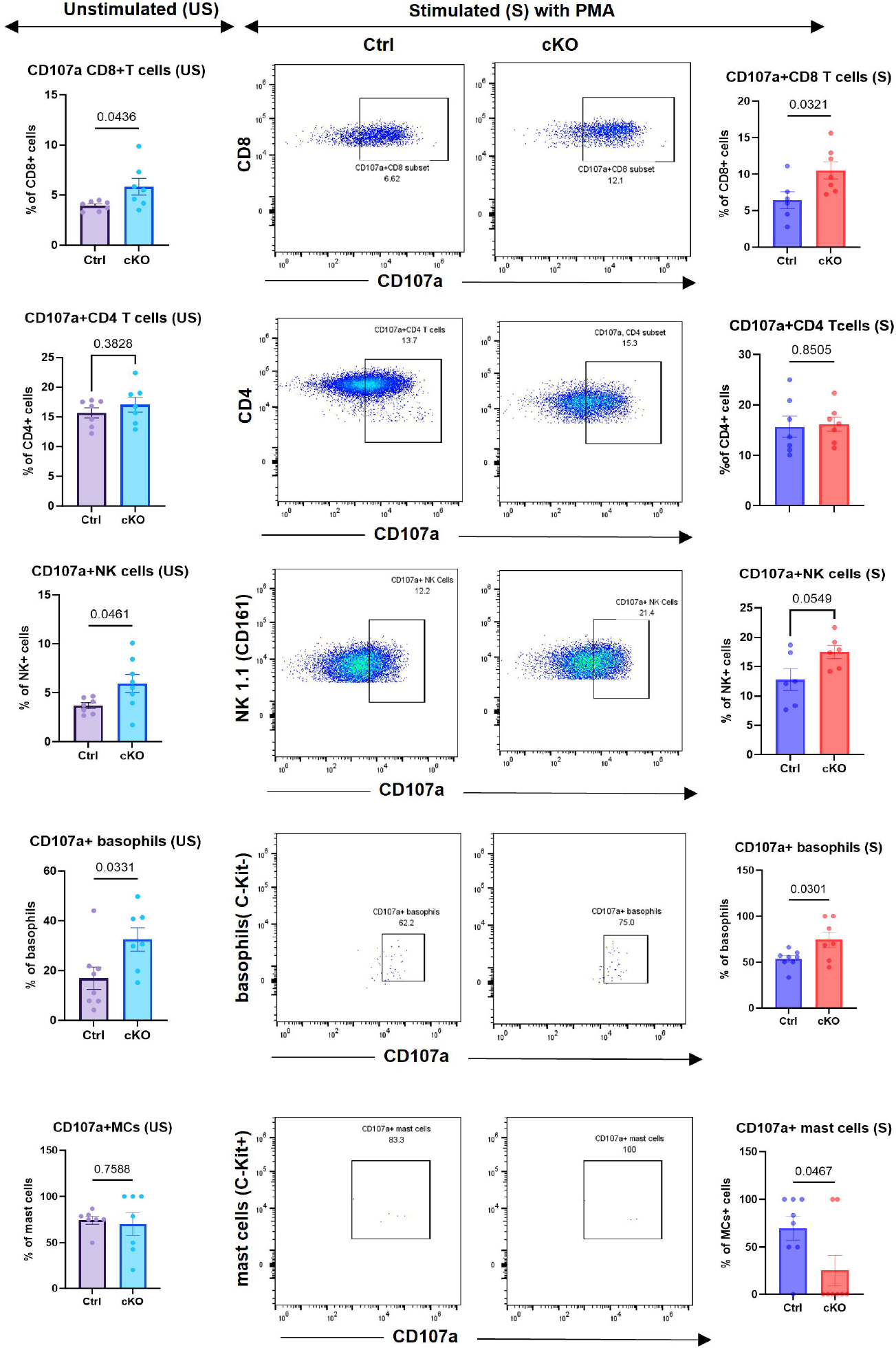
Enhanced immune cell activation in lungs from mice with Cyp11a1-deficient mast cells. CD8^+^ T cells, CD4^+^ T cells, NK cells, basophils and mast cells were assessed for expression of the general activation/degranulation marker CD107a by flow cytometry. Assessments were performed at baseline conditions (unstimulated; US) or after activation with PMA (stimulated; S). The gating strategy is shown. Results are expressed as % CD107a-positve cells out the total cells in the respective population. Data represent mean values ±□SEM; *P*□< □0.05 by unpaired two-tailed *t*-test (at least n = 6/group). US, unstimulated; S, stimulated with PMA.

### Mast cell-derived steroidogenesis restrains T cell effector responses in the tumour microenvironment

The results above indicate that mast cell steroidogenesis impacts on the ability of CD8^+^ T cells to undergo degranulation. Given the central role of T cells in cytotoxicity against tumour cells, we next performed experiments to further assess the effects of mast cell-expressed Cyp11a1 on T cell functionality. To this end, we performed *ex vivo* stimulation of lung immune cells isolated from control and mast cell–specific *Cyp11a1* knockout (cKO) mice. Following PMA activation, both CD4□ helper and CD8□ cytotoxic T cells from cKO mice produced significantly higher levels of interferon-gamma (IFN-γ) compared to those from control animals (**Fig. 7a**). These findings suggest that Cyp11a1-mediated steroid synthesis in mast cells exerts an immunosuppressive influence on T cell activation. In contrast, no significant effect of the depletion of mast cell-expressed Cyp11a1 on IFN-γ expression was seen in NK cells. We also assessed whether mast cell Cyp11a1 has effects on the expression of TNF-α. These analyses revealed that mast cells deficient in *Cyp11a1* had a profoundly reduced ability to express TNF-α protein, whereas no corresponding effects on TNF-α levels were seen in either CD4^+^ T-cells, CD8^+^ T cells, NK cells or B cells (**Fig. 7a**). In agreement with these findings, qPCR analyses confirmed increased expression of IFN-γ mRNA in the lungs of cKO vs control mice, whereas no significant effects of the mast cell-expressed Cyp11a1 on TNF-α mRNA levels were seen (**Fig. 7b**).

**Figure 7.**
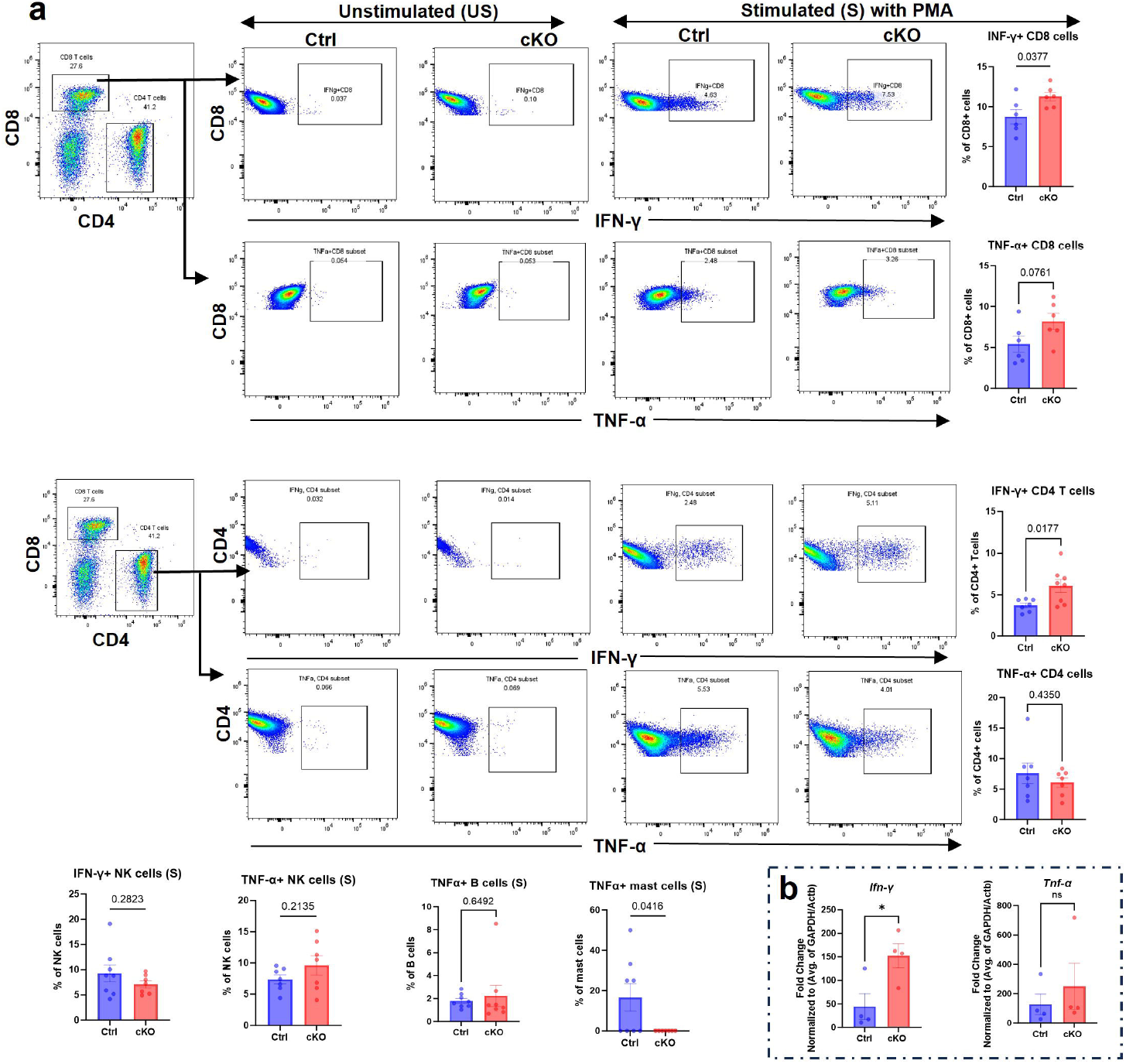
Increased interferon-γ production by T cells in lungs from mice with Cyp11a1-deficient mast cells. Total dispersed lung cell populations were prepared from tumour-bearing control mice (Ctrl) and mice with *Cyp11a1*-deficient mast cells (cKO), and were activated by incubation with PMA. (**a**) After activation of cells, the expression of IFN-γ and TNFα in CD8^+^ T cells, CD4^+^ T cells, NK cells, B cells and mast cells was assessed by flow cytometry as indicated. (**b**) qPCR analysis of IFN-γ and TNFα gene expression in lungs from tumour-bearing control mice (Ctrl) and mice with *Cyp11a1*-deficient mast cells (cKO). Data represent mean□± □SEM; *P*□< □0.05 by unpaired two-tailed *t*-test (at least n = 6/group).

## DISCUSSION

Numerous previous studies have implicated mast cells in melanoma progression. In clinical studies, the presence and abundance of mast cells has frequently been associated with poor prognosis, but there are also clinical studies in which the presence of mast cells has been linked to favourable outcomes [28-33]. The impact of mast cells on melanoma progression has also been studied in animal models. In melanoma models similar to the model used herein, two independent studies have shown that total deficiency in mast cells is associated with a more pronounced lung colonization of melanoma cells, i.e. pointing to an altogether harmful role of mast cells in such settings [34, 35]. In contrast, the growth of the primary tumour was not affected by the total absence of mast cells [34, 35]. Notably, these studies hence reflect the total impact of mast cells in these settings, taking into account the combined effect of all of the compounds that mast cells express under a given condition. As another approach, effects of individual mast cell-expressed compounds can be studied. Intriguingly, such studies have revealed that the absence of mast cell-specific proteases is associated with enhanced lung metastasis and growth of primary tumours in melanoma models [34, 36], pointing to anti-tumour effects mediated by these mast cell-specific enzymes. Hence, this indicates that mast cells can express both tumour-promoting and tumour-suppressive effects, with a conceivable scenario being that the net effect of mast cells on tumour progression reflects the balanced output of such pro- and anti-tumour factors under given tumour conditions.

As noted above, several previous reports have suggested that mast cells may promote melanoma progression. This could potentially be linked to the ability of mast cells to express various tumour-promoting growth factors, including VEGF, FGF and TGF-β [37-39]. However, in this study we have evaluated an alternative scenario by which mast cells may halt melanoma progression. In previous studies it has been shown T cells can be induced to produce steroids in tumour settings, and that such T cell-derived steroids can cause immunosuppression leading to enhanced tumour growth in a melanoma model [26]. Given the reported role of mast cells in regulating melanoma metastasis to the lung [34, 35], we here asked whether the impact of mast cells on melanoma progression can be attributable to mast cell-derived steroid synthesis. This was approached by using mice in which *Cyp11a1* expression was conditionally ablated in mast cells.

In support for a role of mast cell-derived steroids in regulating melanoma metastasis, our data reveal a substantial reduction of metastatic colonisation of the lung in mice in which mast cells lack the expression of Cyp11a1, a key rate-limiting enzyme in steroid synthesis [16, 17]. Intriguingly, mast cell-specific deletion of *Cyp11a1* was associated with an overall decrease in the mast cell numbers in the lungs of tumour-bearing mice, as shown both by staining for mast cells and by analysing the levels of mast cell-specific transcripts. One potential explanation for this finding could be that mast cell-derived steroids act in an autocrine stimulatory effect on mast cell accumulation in the lung. However, the molecular mechanism behind this effect remains to be explored.

Notably, Cyp11a1 was not ubiquitously expressed among the lung mast cell populations, as evidenced by the existence of both *Cyp11a1*^+^ and *Cyp11a1*^-^ mast cells. It was also notable that numerous cells of the lung expressed Cyp11a1 but were negative for mast cell markers. However, the expression of the *Cyp11a1* gene was considerably reduced in lungs of mice with mast cell-specific deletion of Cyp11a1, indicating that mast cells in fact account for a substantial fraction of the total Cyp11a1 expression. The latter was also supported by immunohistochemical staining, which revealed a clear reduction in the overall Cyp11a1 expression in lungs of mice with mast cell-specific deletion of *Cyp11a1*.

An intriguing finding in this study was that the overall positivity for CD107a/LAMP1 was substantially higher in lungs of tumour bearing mice with mast cell-specific deletion of *Cyp11a1* vs control mice. CD107a/LAMP1 is a protein present in the membrane of secretory granules/lysosomes of various immune cell types. In their resting state, CD107a/LAMP1 is hence intracellular, but when the cells are activated, this can lead to degranulation, accompanied by fusion of the granule/lysosome membranes with the cell membrane, accompanied by cell surface exposure of CD107a/LAMP1. Cell surface exposure of CD107a/LAMP1 can thus serve as a marker for immune cell activation/degranulation [40-42]. Hence, the increase in CD107a positivity as a consequence of mast cell-specific deletion of *Cyp11a1* indicates that mast cell-derived steroids may suppress immune cell activation and degranulation. To further substantiate this finding, we employed flow cytometry analysis to evaluate the effects of mast cell Cyp11a1 expression on individual immune cell populations. These analyses did not reveal any significant effects of mast-cell Cyp11a1 on the numbers of either T cells (CD4^+^ or CD8^+^), B cells, NK cells, eosinophils or basophils within the lungs of tumour-bearing mice. However, when assessing the activation status of the respective immune cell populations by assessing CD107a/LAMP1 positivity, we noted that CD8^+^ T cells, NK cells and basophils were activated/degranulated to a higher extent in lungs of mice with mast cell-specific ablation of *Cyp11a1* expression in comparison with controls. Hence, these findings suggest that mast cell-derived steroid synthesis has the capacity to suppress the activation of these respective immune cell populations. Notably, CD8^+^ T cells and NK cells are known to have major roles in the regulation of tumour cell growth [26], and the relieved suppression of these cell populations in the absence of mast cell-derived Cyp11a1 may consequently account, at least partly, for the reduced tumour development seen in these mice.

To provide deeper insight into this issue, we also assessed whether mast cell-specific deletion of *Cyp11a1* had an impact on the expression of classical markers for immune cell activation, by focusing on IFN-β and TNF-α. Indeed, we found that CD8^+^ T cells in lungs from mice with mast cell-specific deletion of *Cyp11a1* exhibited elevated levels of IFN-γ, as assessed by flow cytometry analysis. Elevated IFN-γ levels were also seen in CD4^+^ T cells in lungs from mice with mast cell-specific depletion of *Cyp11a1*.

A possible limitation of this study is that we used only 12-14 weeks-old C57BL/6 mice without control of hair cycle stage, which can influence skin physiology and potentially systemic immune responses. However, the skin is a complex neuroendocrine and immune organ in which local signalling pathways regulate barrier and immune functions [43]. Future studies accounting for hair cycle dynamics or repeating the mice study with a larger cohort size may help to refine these findings. We also did not directly profile steroid metabolites downstream of Cyp11a1. As this enzyme initiates steroidogenesis, multiple bioactive products-including steroid hormones and vitamin D derivatives may contribute to the observed effects [44]. Thus, our findings reflect the impact of pathway disruption rather than specific mediators. More broadly, our results support a role for mast cell-derived steroids in tumor-host communication within the metastatic niche. These findings are consistent with emerging evidence that neuroendocrine pathways shape anti-tumor immunity, suggesting that local steroidogenesis may contribute to systemic immune regulation during metastasis.

Altogether, this study introduces a hitherto unrecognised impact of mast cell-derived steroid synthesis on melanoma metastasis to the lung. These findings may thus be related to the emerging awareness that immunosuppression by steroids can be linked to increased resistance to cancer therapy [45, 46]. Based on the present data, we may envision that targeting of mast cells by regimes that suppress their capacity to release steroids may become explored as potential therapeutic options to suppress melanoma metastasis in clinical settings. However, future work is required to evaluate the impact of our present findings in human clinical settings. It is also important to note that, while we implicate mast cell-derived steroids as modulators of immune suppression, the precise steroid species involved and their downstream signalling pathways remain to be elucidated. Further investigations are also warranted to assess the impact of a mast cell-steroid axis in tumour settings other than melanoma.

## ACKNOWLEDGEMENTS

We would like to thank the Flow Cytometry Facility (Joana Cerveira and Sameen Khan), Confocal Microscopy Facility (Ben Sutcliffe and Jonathan Howe), the Histology Facility at the Department of Pathology, and the UBS Animal Facility (Polly Seaton) at the Gurdon Institute, for technical assistance and animal support. We would like to thank Prof. Heike Laman for support and constructive feedback on the study. We would like to thank Fabio Melo, Uppsala University, for ordering validated primers from the Bio-Rad.

## AUTHOR CONTRIBUTIONS

AA: Planned, designed, and performed experiments; analysed, assembled, visualized, and quantified data; wrote the manuscript.

KB: Assisted with intravenous injections and animal studies.

HH: Assisted with tissue processing, FACS analysis and RNA extraction.

## Figure Legends

**Suppl Fig. 1. Gating strategy used for Figs 5-7**.

**Suppl. Table 1: List of Primers with sequences**

