## Supplementary figures and images for "Mast Cell Specific *Cyp11a1* Deficiency Promotes T Cell Mediated Immunity and Suppresses Tumour Metastasis in a Mouse Model of Melanoma"

### Supplementary Figure 1.jpg

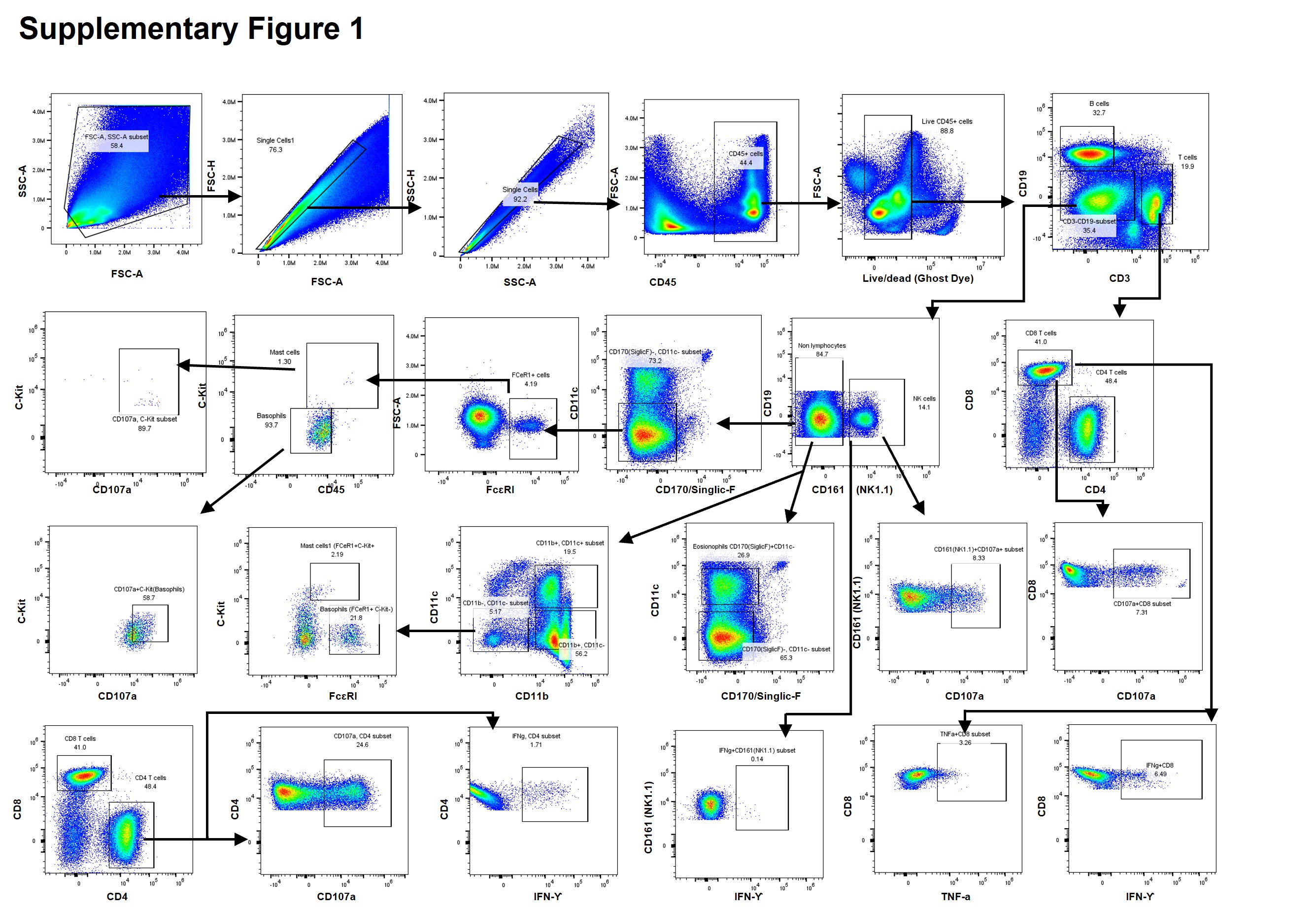

### Supplementary Table 1.jpg

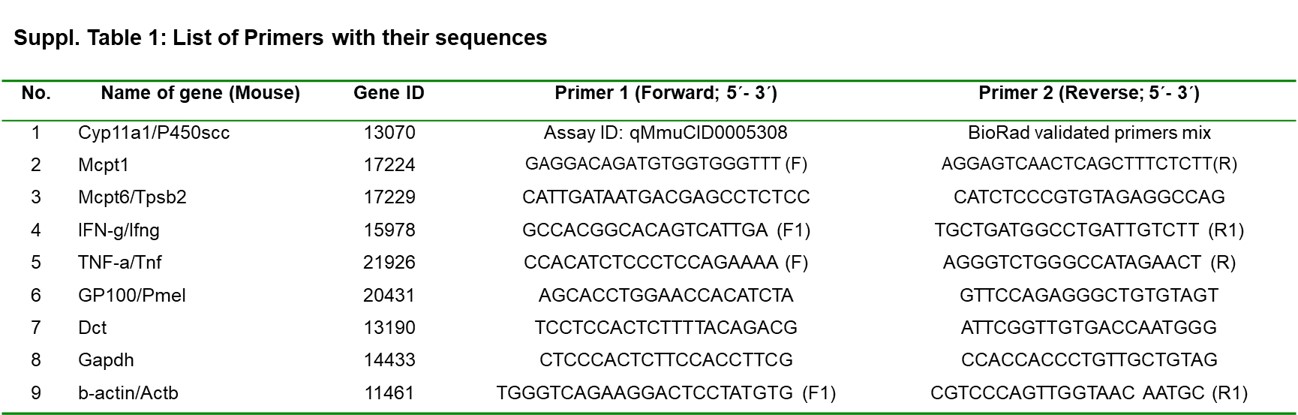
